# CRISPR-Cas9 gene editing and rapid detection of gene-edited mutants using high-resolution melting in the apple scab fungus, *Venturia inaequalis*

**DOI:** 10.1101/2021.02.04.428760

**Authors:** Mercedes Rocafort, Saadiah Arshed, Debbie Hudson, Jaspreet Singh, Joanna K. Bowen, Kim M. Plummer, Rosie E. Bradshaw, Richard D. Johnson, Linda J. Johnson, Carl H. Mesarich

## Abstract

**Background:** Scab, or black spot, caused by the filamentous fungal pathogen *Venturia inaequalis*, is the most economically important disease of apple (*Malus* x *domestica*) worldwide. To develop durable control strategies against this disease, a better understanding of the genetic mechanisms underlying the growth, reproduction, virulence and pathogenicity of *V. inaequalis* is required. A major bottleneck for the genetic characterization of *V. inaequalis* is the inability to easily delete or disrupt genes of interest using homologous recombination. Indeed, no gene deletions or disruptions in *V. inaequalis* have yet been published. Recently, CRISPR-Cas9 has emerged as an efficient tool for gene editing in filamentous fungi. With this in mind, we set out to establish CRISPR-Cas9 as a gene editing tool in *V. inaequalis*.

**Results:** We showed that CRISPR-Cas9 can be used for gene inactivation in the apple scab fungus. As a proof of concept, we targeted the melanin biosynthesis pathway gene *trihydroxynaphthalene reductase* (*THN*), which has previously been shown to result in a light-brown colony phenotype when transcriptionally silenced using RNA interference. Using one of two CRISPR-Cas9 single guide RNAs (sgRNAs) targeted to the *THN* gene, delivered by a single autonomously replicating Golden Gate-compatible plasmid, we were able to identify six of 36 stable transformants with a light-brown phenotype, indicating an ^~^16.7% gene inactivation efficiency. Notably, of these six *THN* mutants, five had an independent mutation. As part of our pipeline, we also report a high-resolution melting (HRM) curve protocol for the rapid detection of CRISPR-Cas9 gene-edited mutants of *V. inaequalis*. This protocol identified a single base pair deletion mutation in a sample containing only 5% mutant genomic DNA, indicating high sensitivity for mutant screening.

**Conclusions:** In establishing CRISPR-Cas9 as a tool for gene editing in *V. inaequalis*, we have provided a strong starting point for studies aiming to decipher the function of genes associated with the growth, reproduction, virulence and pathogenicity of this fungus. The associated HRM curve protocol will enable CRISPR-Cas9 transformants to be screened for gene inactivation in a high-throughput and low-cost manner, which will be particularly powerful in cases where the CRISPR-Cas9-mediated gene inactivation efficiency is low.

## Introduction

Fungal species from the *Venturia* genus are devastating plant pathogens of economically important crops that mainly belong to the *Rosaceae* (1–3). The best researched of these pathogens is *Venturia inaequalis*, which causes scab, or black spot, the most economically important disease of apple (*Malus* x *domestica*) worldwide (1). Under favourable conditions, this disease can result in 70% or more of the crop being lost, as scab renders the apples unmarketable (i.e. through blemishes and deformation), and reduces both the growth and yield of the plant (i.e. by causing repeated defoliation of trees over several seasons) (1, 3, 4). To develop durable control strategies against scab disease, a better understanding of the genetic mechanisms underlying the growth, reproduction, virulence and pathogenicity of *V. inaequalis* is required.

A key development over recent years has been the availability of several *V. inaequalis* genome sequences and gene catalogues (2, 5–9), as well as the development of both polyethylene glycol (PEG)-mediated protoplast and *Agrobacterium tumefaciens*-mediated transformation protocols for use with this fungus (10). However, while several *V. inaequalis* genes of interest that are putatively involved in the infection process of apple have been identified (5, 11–15), none have been functionally characterized to date using traditional gene deletion or disruption techniques. Indeed, no gene deletions or disruptions, based on homologous recombination, have yet been reported for *V. inaequalis* in the literature. This suggests that gene deletion or disruption by traditional homologous recombination is extremely inefficient in *V. inaequalis*. It should be noted that transcriptional silencing of multiple genes in *V. inaequalis* has been achieved using RNA interference (RNAi) (16). However, RNAi does not typically silence a gene to completion and the observed phenotypes can be inconsistent, unclear or absent, making it difficult to determine function (17, 18). Taken together, alternative gene disruption and deletion tools, such as the Clustered Regularly Interspaced Short Palindromic Repeats-Cas9 (CRISPR-Cas9) system (19), are desperately needed to assess gene function in *V. inaequalis*.

The CRISPR-Cas9 system is a powerful tool for gene editing that has been established in many species of filamentous plant-pathogenic microbes. Indeed, CRISPR-Cas9 has been used to generate gene inactivations in more than 40 species of filamentous fungi and oomycetes (20), including *Phytophthora sojae* (21), *Magnaporthe oryzae* (22), *Ustilago maydis* (23) and a fungus that is closely related to *V. inaequalis, Leptosphaeria maculans* (24).

The CRISPR-Cas9 gene editing system requires two components: the Cas9 endonuclease and a single guide RNA (sgRNA). The Cas9 endonuclease is an RNA-guided enzyme that generates a double strand break (DSB) in the genome. The sgRNA consists of a protospacer sequence of 20 nucleotides at the 5’ end that targets specific DNA by base pairing, and an 80-nucleotide scaffold structure that binds to Cas9. The sgRNA-Cas9 complex only cleaves the target DNA if it is flanked by a protospacer motif (PAM) (19).

After the Cas9 endonuclease generates a DSB in the target DNA, DNA repair mechanisms are activated (19, 25). The DNA can be repaired by a non-homologous end-joining mechanism (NHEJ) or homology-directed repair (HDR), although NHEJ is usually the dominant DNA repair pathway in fungi (26). DNA repair by NHEJ is error-prone and is likely to introduce small insertions/deletions (indels) or nucleotide substitutions, which can lead to frameshift mutations that cause a gene disruption (25). Alternatively, a double-stranded DNA template (donor DNA), usually harbouring a selectable marker, can be introduced to use as a repair template for HDR.

Traditionally, screening for the identification of mutants generated by the CRISPR-Cas9 system can be achieved using an enzymatic mismatch cleavage (EMC) method (27) or a polyacrylamide gel electrophoresis (PAGE)-based method (28) that both rely on the detection of DNA heteroduplexes. Both EMC and PAGE detect large indels with a similar efficiency; however, the sensitivity with which they detect small indels (i.e. the type of indels that are usually generated by CRISPR-Cas9 DSB) is low (27–29). Alternatively, CRISPR-Cas9 mutants can be identified by amplicon sequencing, which is a tedious and expensive process. High-resolution melting (HRM) curve analysis is a fluorescence-based technique that measures the melting temperature of double-stranded DNA and, in doing so, can discriminate between amplicons with different melting temperatures (30–32). HRM curve analysis has been widely used to identify mutations and single nucleotide polymorphisms in various genes (30, 33), and has recently been used to reliably identify CRISPR-Cas9-mediated base pair (bp) indels in plants (29, 34). To date, even though HRM curve analyses are largely used for fungal species identification, to our knowledge, an HRM curve analysis has not yet been employed to screen for CRISPR-Cas9-generated mutants in fungi.

In this study, we set out to establish the CRISPR-Cas9 gene editing system in *V. inaequalis*. For this purpose, we first generated the Golden Gate-compatible plasmid Cas9HygAMAccdB by modifying the previously published Cas9 autonomously replicating plasmid ANEp8_Cas9_LIC1 (35). Here, Golden Gate-compatibility was chosen as it enabled the introduction of a sgRNA into Cas9HygAMAccdB using a single step, facilitating the creation of a Cas9HygAMA-*sgRNA* plasmid in less than one week. Next, using the melanin biosynthesis pathway gene *trihydroxynaphthalene reductase* (*THN*) as a proof of concept, we showed, in conjunction with the Cas9HygAMA*-sgRNA* plasmid, that the CRISPR-Cas9 gene editing system can be successfully applied to *V. inaequalis*. As part of this process, we also established a method based on an HRM curve analysis for the high-throughput screening of CRISPR-Cas9 gene-edited mutants of *V. inaequalis*.

## Results and discussion

### CRISPR-Cas9 can be used for gene disruption in *V. inaequalis*

We set out to establish the CRISPR-Cas9 gene editing system in *V. inaequalis*, using the melanin biosynthesis pathway gene *THN* (Joint Genome Institute ID: *atg4736.t1*) as a target for inactivation. The *THN* gene was chosen as a proof of concept, as a previous study had shown that *V. inaequalis* displays a distinctive light-brown phenotype when this gene is transcriptionally silenced using RNAi, indicative of reduced melanisation (16). In this way, transformants of *V. inaequalis* inactivated for the *THN* gene using the CRISPR-Cas9 gene editing system can be rapidly identified through a simple visual screen. For ease of use, we chose to employ a CRISPR-Cas9 gene editing system that, similar to the one previously established in *Aspergillus niger* (35), only requires a single autonomously replicating plasmid, Cas9HygAMAccdB (containing both the Cas9 endonuclease and sgRNA), for gene inactivation.

The Cas9HygAMAccdB plasmid contains an *A. niger* codon-optimized *cas9* gene expressed under the control of the *pkiA* (*pyruvate kinase*) promoter. To ensure expression in the fungal nucleus, the Cas9 endonuclease was tagged at its carboxyl (C) terminus with a nuclear localization signal (NLS). The Cas9HygAMAccdB plasmid also contains an RNA polymerase III promoter to facilitate expression of the sgRNA *in vivo*. Using the chosen CRISPR-Cas9 system, the 20 nucleotides of the sgRNA protospacer were synthesized as two pairs of complementary oligonucleotides that were pre-annealed and cloned into Cas9HygAMAccdB by a single-step Golden Gate reaction, enabling Polymerase Chain Reaction (PCR)-free cloning that could be completed in less than one week.

An autonomously replicating plasmid was chosen as it has several advantages. Firstly, autonomously replicating plasmids can enhance fungal transformation efficiency, as recombination between the plasmid and chromosome is not required (36). As such, autonomously replicating plasmids can be used in fungal species that exhibit low transformation efficiency, such as *V. inaequalis*. Secondly, autonomously replicating plasmids could be lost once selection (e.g., as mediated through hygromycin B) is removed (36). In doing so, autonomously replicating plasmids can reduce off-target effects by only enabling transient expression of the Cas9 endonuclease in the fungus (37). As such, autonomously replicating plasmids could be recycled, which would enable the sequential inactivation of genes in CRISPR-Cas9 gene-edited mutants and could also facilitate the subsequent complementation of mutants generated using CRISPR-Cas9 technology.

As a starting point for inactivation, sgRNAs were designed to target the amino (N) terminus (first and second exon) instead of the C terminus of the *THN* gene. This is because mutations at the C terminus of a gene are less likely to cause a frameshift mutation that results in inactivation (38). As different sgRNAs can display different targeting efficiencies (38), two different sgRNAs, sgRNA 4 and sgRNA 20, with similar predicted on-target activity, and no predicted off-target activity (Table 1), were selected for inactivation of the *THN* gene. Special attention was taken to ensure that the sgRNAs did not target any other melanin biosynthesis pathway genes that have a high degree of conservation to *THN*, such as the *1,3,6,8-tetrahydroxynaphthalene reductase* gene (Joint Genome Institute ID: *atg3631.t1*). PEG-mediated protoplast transformation of *V. inaequalis* with sgRNA 4, targeting the first *THN* exon (Figure 1.A), resulted in 98 independent transformants, of which 62 ceased to grow on hygromycin B selection media after one week, and were therefore considered transient, giving a final number of 36 stable transformants. Transient transformants have been reported in a large number of PEG-mediated protoplast transformations of fungi (39, 40), and it has previously been reported that up to 98% of PEG-mediated protoplast transformants of *V. inaequalis* are transient (10). Of course, it remains possible that the large number of transient transformants generated in our study was the result of premature loss of the autonomously replicating plasmid used to deliver the sgRNA and Cas9.

**Table 1:**
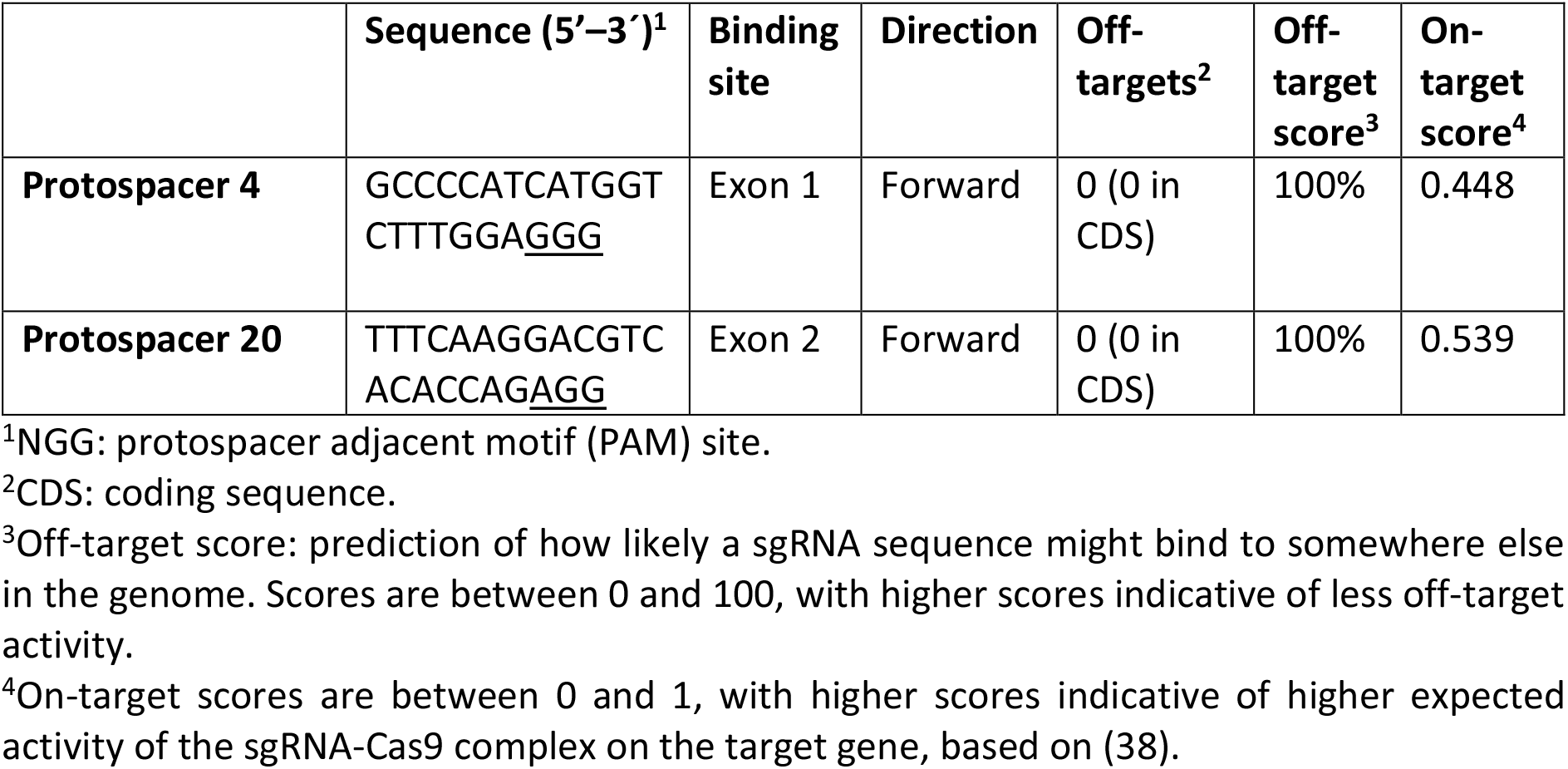
Selected sgRNA protospacers used to target the *Venturia inaequalis* melanin biosynthesis pathway gene *trihydroxynaphthalene reductase* (*THN*).

**Figure 1.**
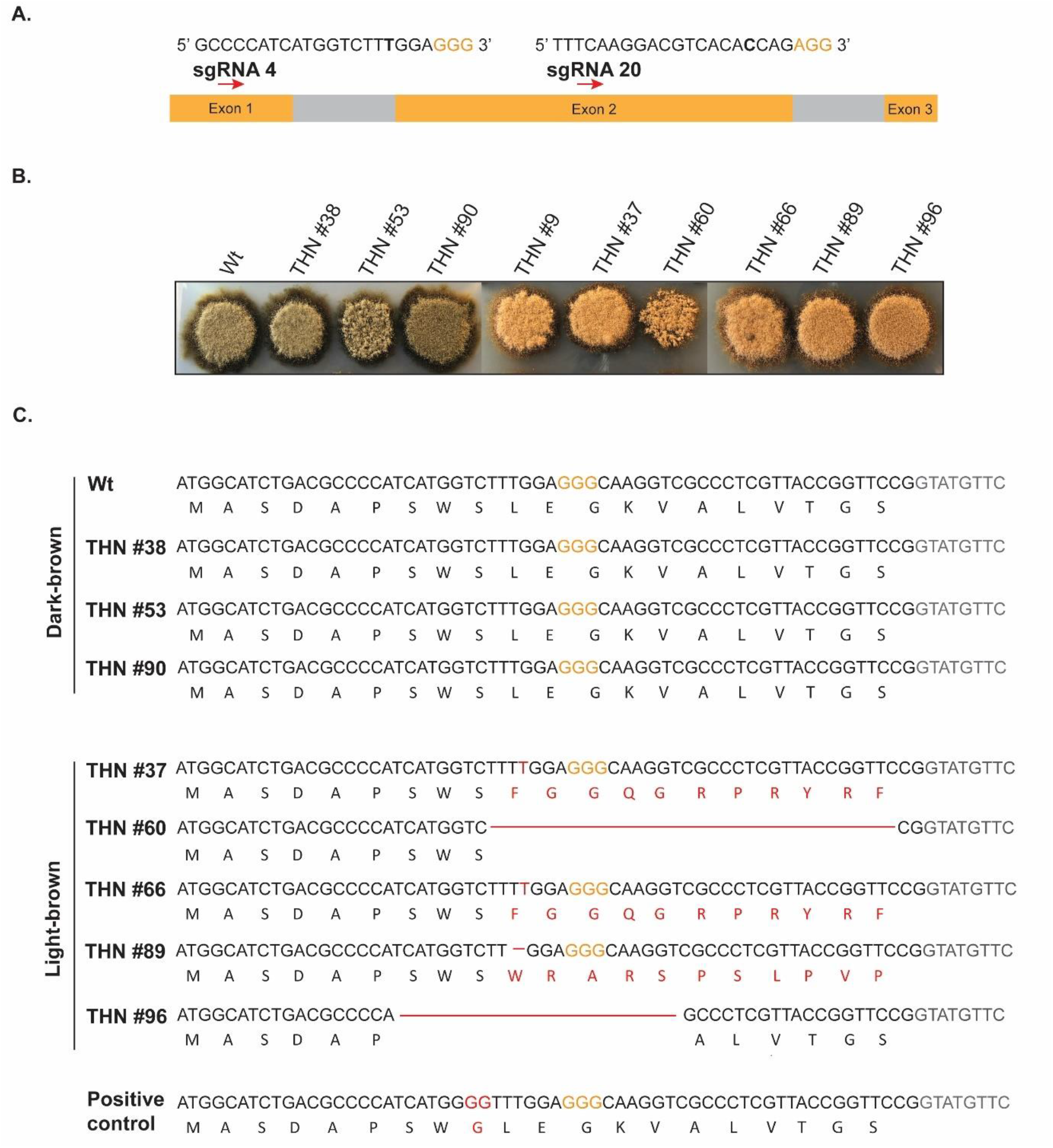
Establishment of the CRISPR-Cas9 gene editing system in *Venturia inaequalis*. **A**. Schematic representation of the *V. inaequalis* melanin biosynthesis pathway gene *trihydroxynaphthalene reductase* (*THN*; 911 bp) with binding sites for the two selected sgRNAs used in CRISPR-Cas9 gene editing experiments shown. Orange: gene exons; Grey: gene introns; Bold nucleotides: expected sgRNA cleavage site; orange nucleotides: PAM site. Arrows: binding direction of sgRNAs. **B.** Colony phenotype of wild type (wt) *V. inaequalis*, and three dark-brown and six light-brown CRISPR-Cas9 transformants grown on potato-dextrose agar at 22°C for 14 days. **C.** Spectrum of CRISPR-Cas9-generated mutations in the *THN* gene. Black nucleotides: exon; Grey nucleotides: intron; Orange nucleotides: PAM site; Red amino acids/nucleotides: mutations observed. Light-brown mutant THN #9 was not sequenced due to a lack of PCR amplification for the *THN* gene.

Notably, of the 36 stable transformants, six had a light-brown phenotype (Figure 1.B). These were transformants THN #9, THN #37, THN #60, THN #66, THN #89 and THN #96. To validate the presence of a mutation in the *THN* gene, it was amplified from each of the six putative mutants by PCR, as well as from three dark-brown transformants without the light-brown phenotype (THN #38, THN #53 and THN #90), and subjected to amplicon sequencing. As expected, all three dark-brown transformants did not contain a mutation in their *THN* gene (Figure 1.C). For the putative light-brown mutant THN #9, the *THN* gene could not be amplified by PCR using two different sets of primers (MR161-MR162, MR185-MR186). This was despite the fact that both *THN*-flanking genes could be amplified, suggesting that a large deletion at the *THN* locus might have occurred (Figure S1). In contrast to the dark-brown transformants, the remaining light-brown *THN* transformants displayed a range of mutations. More specifically, these were a single bp insertion (T) at the same location in THN #37 and THN #66, a 33-bp deletion (TTTGGAGGGCAAGGTCGCCCTCGTTACCGGTTC) in THN #60, a single bp deletion (T) in THN #89, and a 24-bp deletion (TCATGGTCTTTGGAGGGCAAGGTC) in THN #96. Therefore, from the 36 stable transformants, six had a confirmed mutation, giving a gene inactivation efficiency of ~16.7%.

CRISPR-Cas9 gene inactivation efficiencies in other filamentous fungi, in experiments that rely on NHEJ, range between 10 and 100% (22, 23, 41–43). In cases where the CRISPR-Cas9 NHEJ-based gene inactivation efficiency is low, the gene inactivation efficiency could be improved greatly by the incorporation of a donor DNA that is integrated into the genome using HDR (22). Nevertheless, gene inactivation efficiencies between experiments cannot be compared, as these efficiencies will greatly depend on Cas9 expression, sgRNA design and accessibility of the target gene, among other factors (20). Therefore, even though the inactivation efficiency of the *V. inaequalis THN* gene with sgRNA 4 is at the low end, it is likely to vary between genes and no conclusions can yet be drawn as to the overall efficiency of the technique.

Remarkably, PEG-mediated protoplast transformation of *V. inaequalis* with sgRNA 20, targeting the second exon of the *THN* gene (Figure 1.A), resulted in a similar number of stable independent transformants on hygromycin B selection media (31 in total), but none of these transformants had the distinctive light-brown phenotype. This stark difference in the number of transformants inactivated for the *THN* gene is interesting, given that both sgRNAs had a similar predicted on-target activity score (Table 1). However, different sgRNAs can vary greatly in efficiency, as with that previously observed for the *yA* gene of the filamentous fungus *A. nidulans* (44), highlighting the importance of reliable methods to estimate sgRNA efficiency. Thus, while genes will differ in their ability to be inactivated using the CRISPR-Cas9 gene editing system (e.g. due to their location in the genome) (20), it is important that future studies consider multiple sgRNAs for successful gene inactivation in *V. inaequalis*.

### The autonomously replicating CRISPR-Cas9 gene editing plasmid is rapidly lost in most transformants once selection is removed

With the finding that the CRISPR-Cas9 gene editing system could be successfully applied to *V. inaequalis*, we next set out to determine whether this fungus can lose the autonomously replicating plasmid, Cas9HygAMA-*sgRNA*, once the hygromycin B selection is removed. For this purpose, all six light-brown *THN* mutants (THN #9, THN #37, THN #60, THN #66, THN #89 and THN #96), as well as three dark-brown transformants (THN #38, THN #53 and THN #90), all derived from the transformation of *V. inaequalis* with sgRNA 4, were single-spore purified and replica-plated onto both potato-dextrose agar (PDA) and PDA supplemented with 50 μg/ml hygromycin B. After only one round of single-spore isolation and sub-culturing, four of the *THN* mutants (THN #60, THN #66, THN #89 and THN #96) and two of the transformants without the light-brown phenotype (THN #53 and THN #90) lost the Cas9HygAMA-*sgRNA* plasmid and were therefore unable to grow on PDA supplemented with hygromycin B, indicating loss of the Cas9HygAMA-*sgRNA* plasmid (Figure S2).

### High-resolution melting analysis is a sensitive and high-throughput method to screen for CRISPR-Cas9 mutants

In our study, mutants of *V. inaequalis* with a CRISPR-Cas9-mediated gene inactivation of the *THN* gene could be rapidly identified based on their light-brown colony phenotype. However, not all genes of *V. inaequalis* will result in an observable phenotype when inactivated or mutated. For this reason, a low-cost, high-throughput method is required to rapidly identify CRISPR-Cas9 gene-edited mutants of *V. inaequalis* that lack an observable phenotype on a transformation plate facilitating selection. One such method is an HRM curve analysis, which enables the rapid identification of single bp indels in DNA amplicons from transformant genomic DNA (30–32). We set out to test the efficiency of an HRM curve analysis for the detection of *V. inaequalis THN* mutants generated using CRISPR-Cas9 sgRNA 4. As a starting point for this analysis, we first generated a positive control sequence for detecting indels relative to the wild type (wt) sequence (Figure 1.C). The positive control was generated by introducing a two-nucleotide substitution into a PCR amplicon of the *THN* gene using site-directed mutagenesis.

According to the literature, an HRM curve analysis can be affected by different parameters such as genomic DNA quality, PCR amplicon size, amplicon GC content and fluorescent dye used (45). To ensure reliability of the assay, good quality genomic DNA should be extracted. Likewise, the same DNA preparation method should be used across all samples to ensure uniformity in genomic DNA quality. Amplicon size is another crucial parameter that affects the HRM curve analysis. Therefore, two different primer sets were designed to generate amplicons of 230 bp (MR170-MR171) and 123 bp (MR172-MR173) (Table 2). The HRM curve assay was performed with both primers sets using wt genomic DNA and the engineered positive control as DNA template. Specific DNA amplicons could be amplified using both primers sets (Figure 2.A); however, the smaller amplicon showed a clearer shift in the melting curve between wt and positive control (Figure 2.B). Given that smaller amplicons are more suitable for HRM curve analysis (45), and because the smaller amplicon showed a clearer shift in the melting curve, we decided to use the 123 bp amplicon for our screen (primer set MR172-MR173).

**Table 2.**
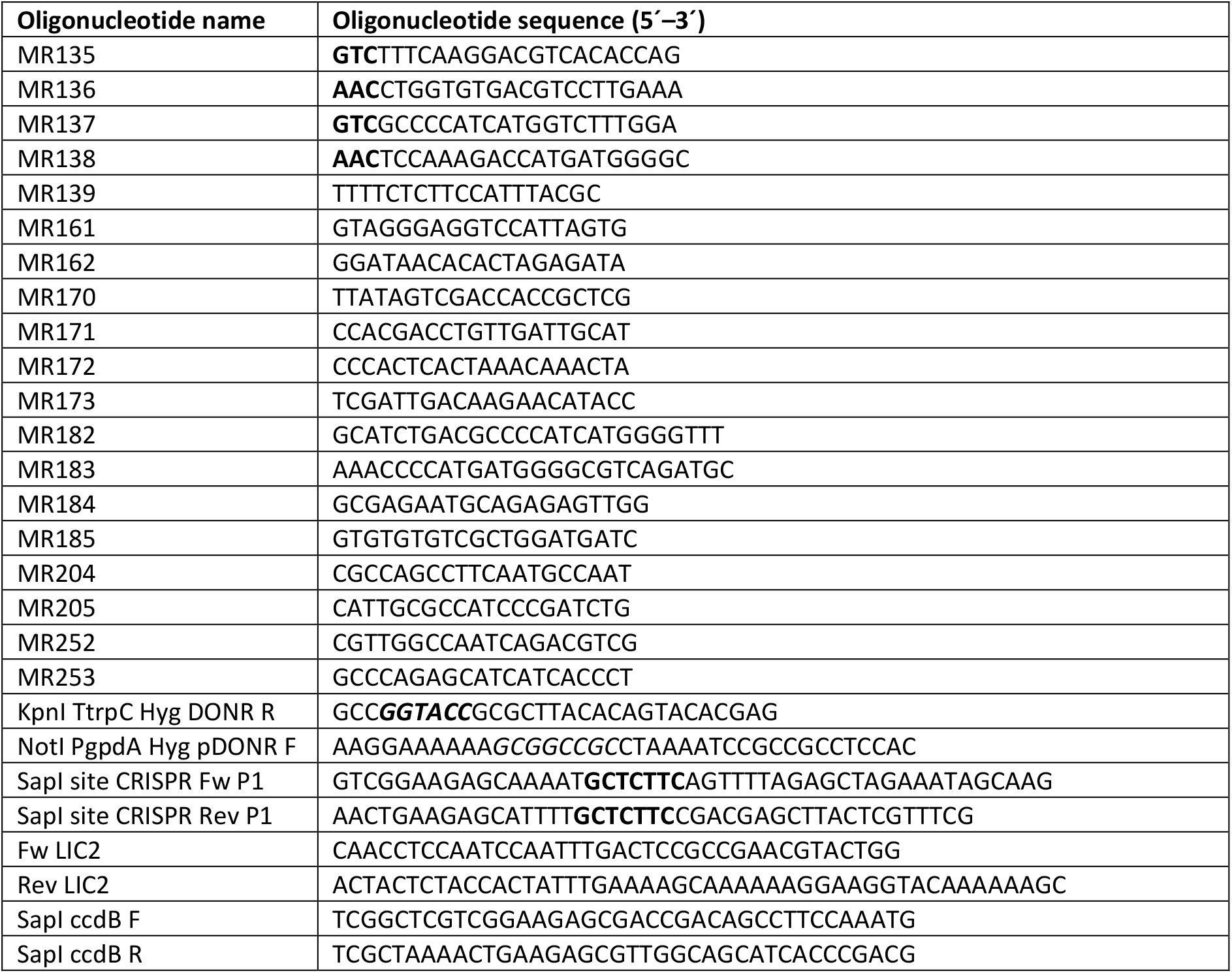
Oligonucleotides used in this study. Bold sequence corresponds to the *Sap*I restriction site. Italicised sequence corresponds to the *Not*I restriction site. Bold italicized sequence corresponds to the *Kpn*I restriction site.

**Figure 2.**
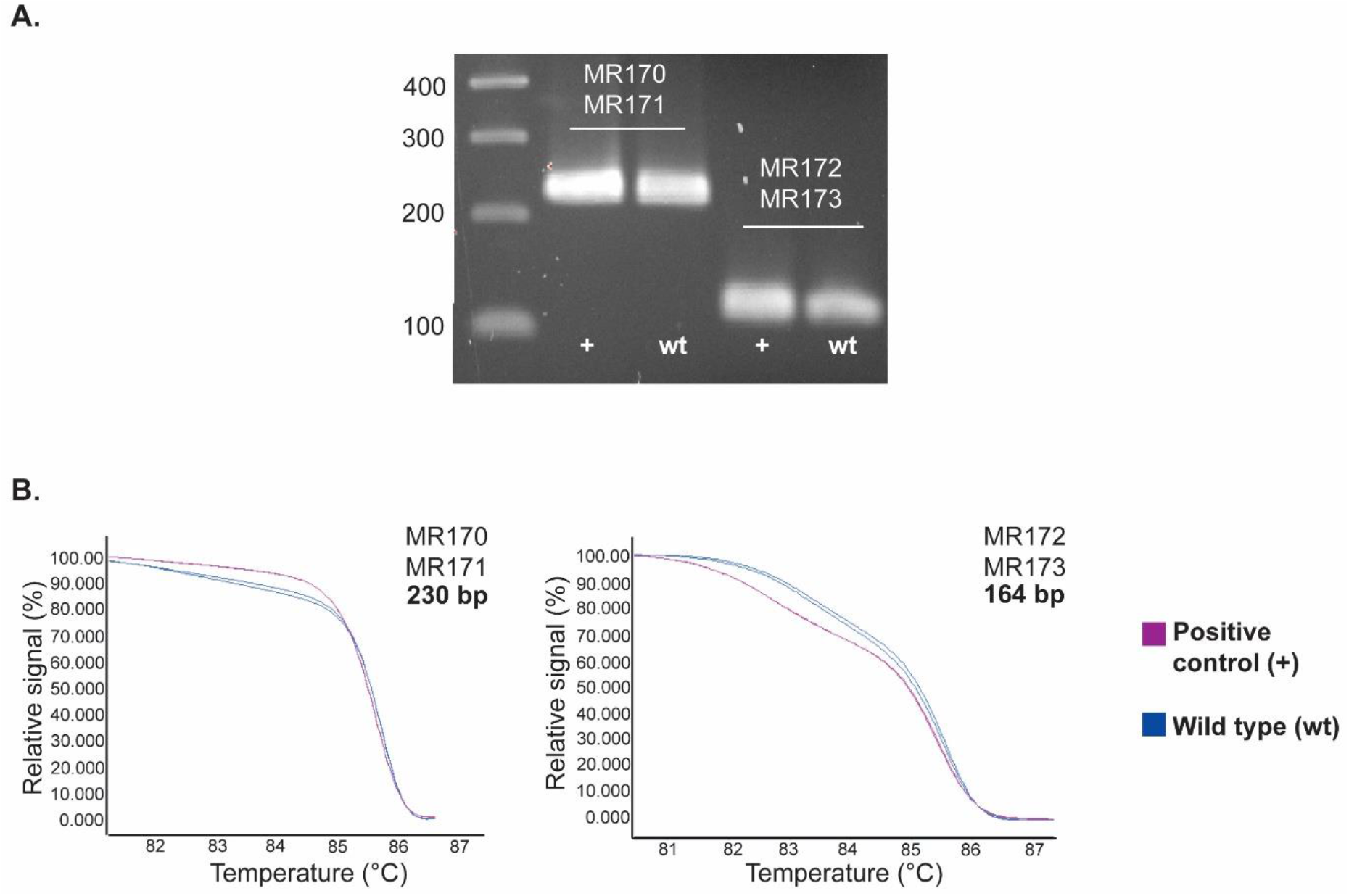
Optimization of primers for qPCR-HRM curve analysis of the *trihydroxynaphthalene reductase* (*THN*) gene mutants generated using CRISPR-Cas9 sgRNA 4. **A.** Conventional PCR amplification of the *THN* gene positive control (+) and wild type (wt) sequences with the different primer sets resolved by electrophoresis on a 1.5% TBE agarose gel. Ladder sizes are shown in base pairs (bp). **B.** Normalized and shifted melting curves of the *THN* gene positive control (+) and wild type (wt) PCR amplicons generated with two different primer sets (MR170-MR171 and MR172-MR173).

Initially, the Bioline SensiFAST^™^ SYBR^®^ No-ROX Kit fluorescent dye was tested to perform the HRM curve analysis; however, no differences between the wt and engineered positive control DNA could be detected (data not shown). It is known that the fluorescent dye used for an HRM curve analysis is crucial to ensure good assay sensitivity, and that high-saturating dyes designed to increase sensitivity are commercially available. With this in mind, the HRM was repeated using the AccuMelt HRM SuperMix high saturating dye SYTO 9^TM^, resulting in clear separation of unique melting curves between mutants (Figure 3). The normalized HRM curves showed that amplicons from wt fungus, as well as dark-brown transformants, clustered together into a group with a similar melting curve profile (blue curves in Figure 3). In contrast, amplicons from the engineered positive control (green curves in Figure 3) and the light-brown *THN* mutants showed distinct melting curves (green, red, grey and purple curves in Figure 3) that correlated with the different mutations seen in Figure 1.C. These results indicate that the HRM curve assay can not only efficiently identify CRISPR-Cas9 gene-edited mutants, but can also discriminate between different mutations. The light-brown mutant THN #9 could not be screened using the HRM curve analysis due to a lack of PCR amplification for the *THN* gene, suggesting that a large deletion at the *THN* locus has been generated. This highlights one of the main limitations of the HRM curve analysis, in that it is not suitable for the detection of large deletion mutants. However, given that such deletion mutants can be easily assessed using standard PCR, this is not an issue.

**Figure 3.**
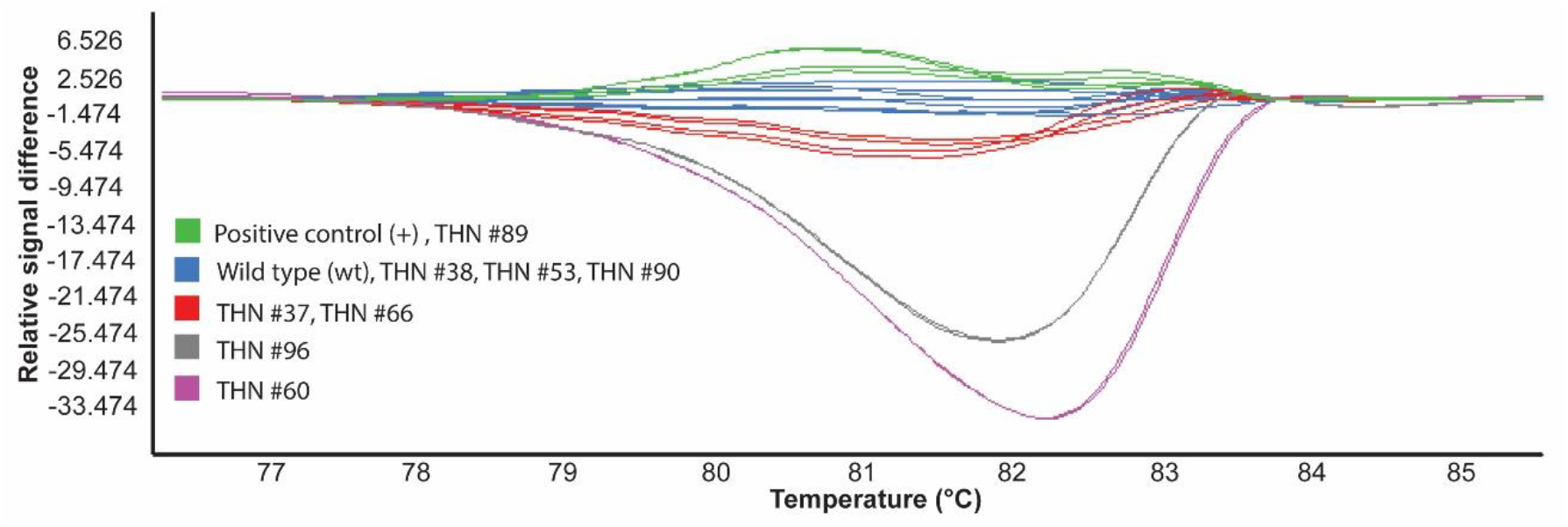
Screening of CRISPR-Cas9 transformants of *Venturia inaequalis* using qPCR-HRM curve and PCR amplicon sequencing analyses. Plot showing normalized and temperature-shifted differences in melting curves between *trihydroxynaphthalene reductase* (*THN*) amplicons of transformants. Melting curve groups were generated by LightCycler^®^ 480 gene scanning software with a sensitivity of 0.40 and using AccuMelt HRM SuperMix fluorescent dye (DNAture). The experiment is based on two technical replicates per sample.

One of the main advantages of the HRM curve analysis is that it can detect mutant DNA that is mixed in with wt DNA, even when the amount of mutant DNA is as low as 1–5% (29, 34). To test the efficiency of the HRM curve assay, genomic DNA from one mutant with a single bp insertion (THN #66) was mixed with wt genomic DNA at different mutant:wt ratios (1:99, 5:95, 10:90, 20:80, 30:70 and 50:50), similarly to that performed by (29). Under our conditions, the assay could not differentiate mutant and wt in 1:99 mutant:wt ratio; however, the mutant DNA could be identified by the HRM curve analysis in all of the other ratios, with a clear shift in the melting curve, indicating an assay sensitivity as high as 5% (Figure 4). Therefore, mutants do not need to be single-spore purified before performing an HRM curve analysis, and multiple mutants can be pooled together for large-scale screens. The observation that mutant DNA can be detected when mixed with wt DNA further reduces the cost and workload of the assay, and further validates the use of an HRM curve analysis for the high throughput screening of CRISPR-Cas9 fungal transformants.

**Figure 4.**
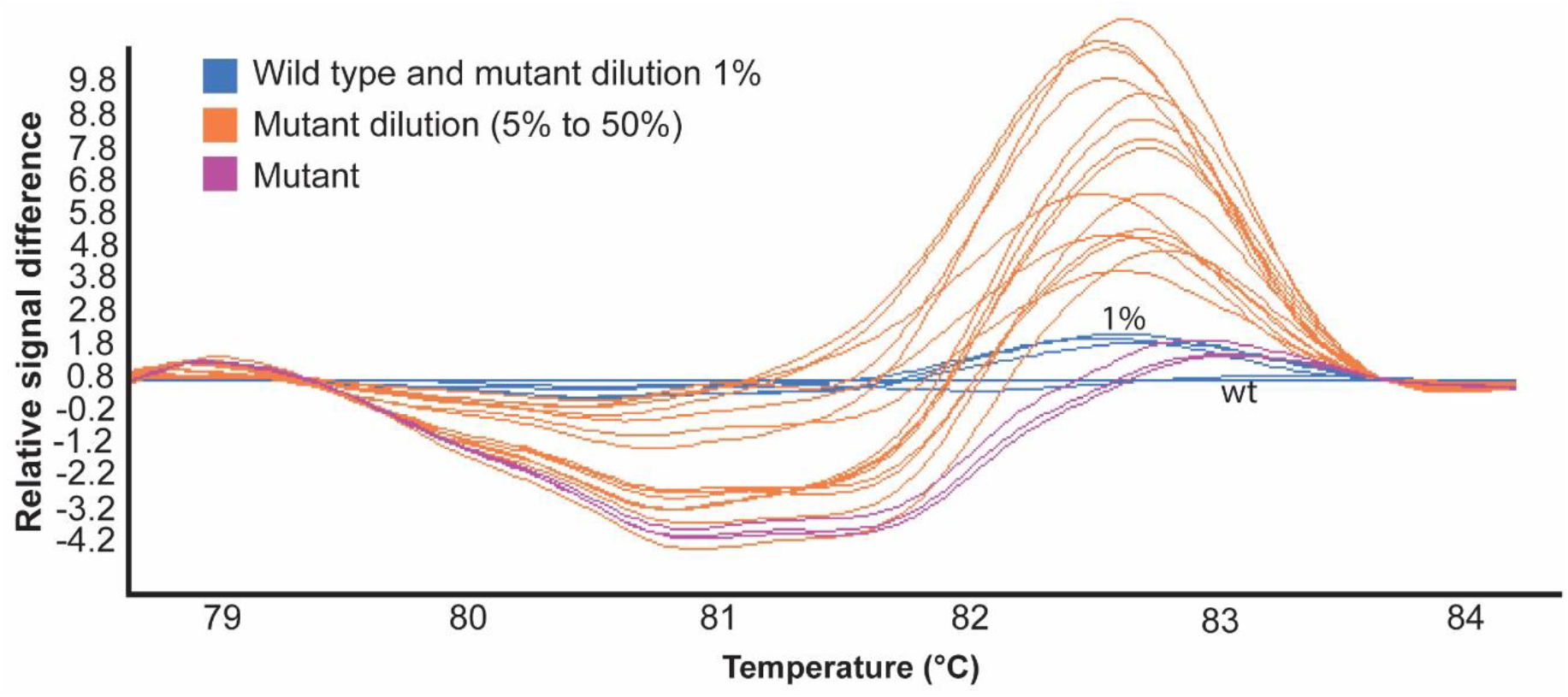
qPCR-HRM sensitivity to detect a one base pair deletion in mutant–wild type DNA mixtures. Plot showing normalized and temperature-shifted differences in melting curves between wild type (wt) *trihydroxynaphthalene reductase* (*THN*) DNA samples of *V. inaequalis* wt mixed with mutant *THN* DNA samples to detect the resolution limit of the HRM curve assay. Melting curve groups generated by LightCycler^®^ 480 gene scanning software with a sensitivity of 0.48 and using AccuMelt HRM SuperMix fluorescent dye (DNAture). A minimum of two technical replicates were performed per sample. Wt genomic DNA and mutant THN #66 genomic DNA were diluted in different ratios (mutant:wt): 1:99, 5:95, 10:90, 20:80, 30:70 and 50:50.

### CRISPR-Cas9 editing of the *THN* gene does not alter the phenotype of *V. inaequalis* grown in culture

CRISPR-Cas9-based experiments can sometimes be detrimental to the target organism due to Cas9 toxicity and/or off-target mutations (46), even though CRISPR-Cas9 off-target mutations have been reported to be unlikely in different filamentous fungi (22, 23, 47). To test if CRISPR-Cas9-mediated transformation has greatly affected the phenotype of *V. inaequalis* (e.g. through toxicity or off-target effects), we investigated the phenotypes of the wt, three dark-brown transformants (THN #38, THN #53, THN #90) and six light-brown mutants (THN #9, THN#37, THN#60, THN#66, THN#89, THN#96) of *V. inaequalis* on and in cellophane membranes overlaying PDA. During growth in cellophane membranes, *V. inaequalis* undergoes morphological differentiation, similar to that observed under the cuticle *in planta*, where it develops infection structures called runner hyphae and stromata (12). Therefore, we investigated whether the *THN* mutants maintained their ability to develop runner hyphae and stromata in cellophane membranes after CRISPR-Cas9-mediated transformation (Figure 5). All mutants showed a similar phenotype to wt, in that they were able to penetrate the cellophane membrane to develop runner hyphae and stromata. Likewise, all mutants maintained their ability to sporulate on the cellophane membrane surface. Taken together, these results suggest that CRISPR-Cas9-mediated transformation has not greatly affected the phenotype of *V. inaequalis*. A more in-depth analysis based on whole genome sequencing is the next step to determine whether this experiment has resulted in any off-target mutations.

**Figure 5.**
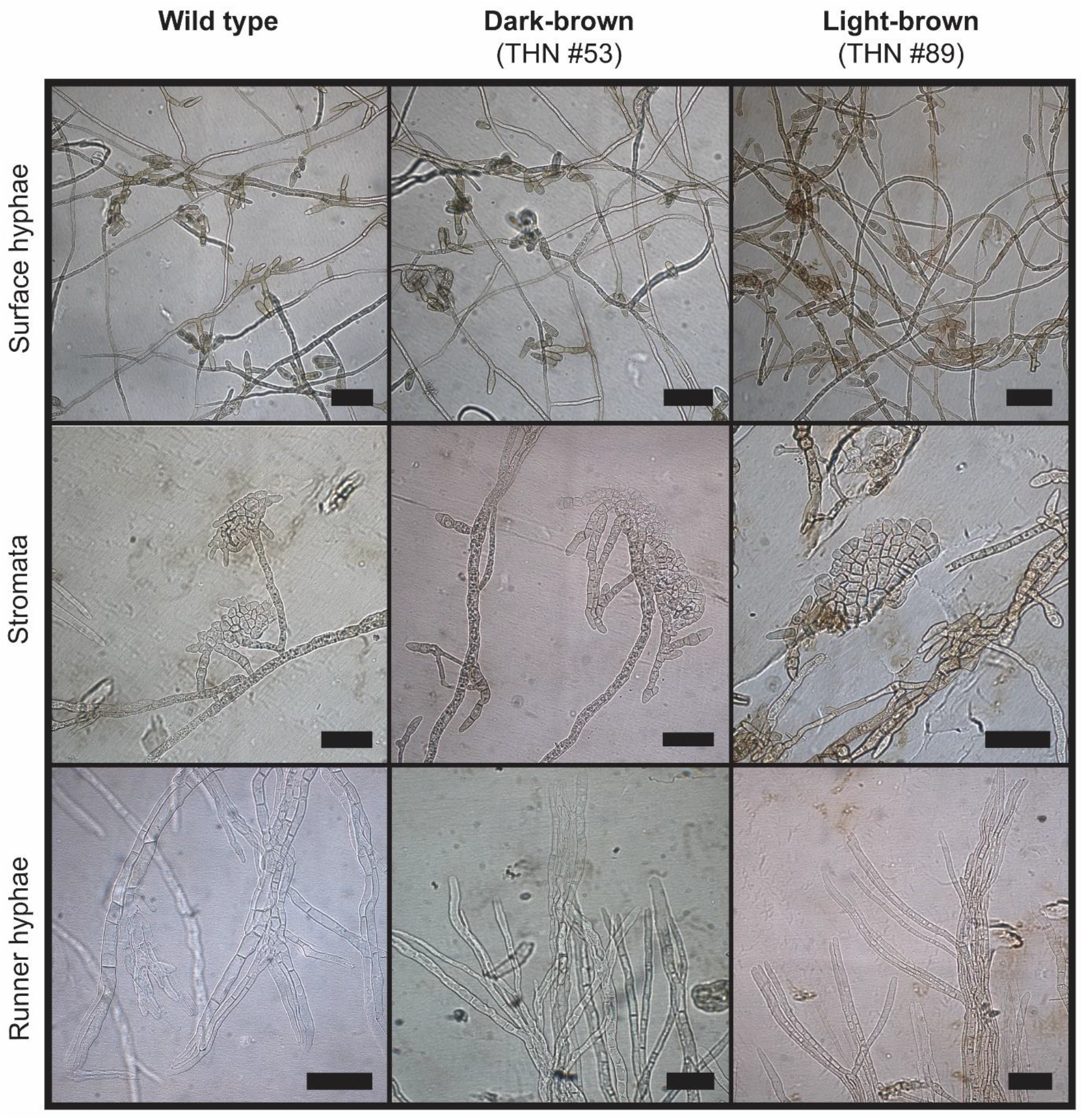
In-culture phenotype of wild-type and CRISPR-Cas9 transformants of *Venturia inaequalis* on and in a cellophane membrane. The conidia of each strain were plated on a cellophane membrane overlaying potato-dextrose agar, followed by incubation at 22°C for 10 days. Scale bar 50 μM. Pictures are representative of all *THN* mutants identified.

### Conclusions

We have successfully applied CRISPR-Cas9 gene editing to the filamentous fungal pathogen, *V. inaequalis*, providing an opportunity for future studies to characterise gene functions associated with the growth, reproduction, virulence and pathogenicity of this fungus. Given that the genomes of several other species from the *Venturia* genus have recently been sequenced (5, 48–53), this development will likely be useful for the functional characterization of genes in these species. Notably, genome sequencing has revealed that members of the *Venturia* genus contain large expanded families of putative effector genes that likely play an important role in host colonization (5). As the functional characterization of gene families is often hindered by functional redundancy between family members, and because the sequential deletion of family members using standard homologous recombination is limited by the number of selectable marker genes that are available, our finding that CRISPR-Cas9 technology can be applied to *V. inaequalis* is also expected to greatly facilitate the functional characterization of effector gene families in the *Venturia* genus (i.e. through sequential gene deletion/disruption or simultaneous gene targeting (multiplexing)) (54).

In addition to applying CRISPR-Cas9 gene editing to *V. inaequalis*, we have developed a high-throughput screening protocol based on an HRM curve analysis for the identification of CRISPR-Cas9-generated mutants of this fungus with as little as one bp insertion or deletion. We have observed that this highly sensitive method can detect mutant DNA even when mixed in a 5:95 mutant:wt ratio, making it an excellent method for high-throughput screening, as mutants do not need to be single-spore purified prior to screening. This method will be of great value for the identification of mutants generated by CRISPR-Cas9 technology in different fungal species where the mutation efficiency is low.

## Methods

### Strains used and growth conditions

*V. inaequalis* isolate MNH120 from New Zealand (ICMP 13258; (55)) was used for CRISPR-Cas9 experiments, and was grown on a cellophane membrane (Waugh Rubber Bands) overlaying PDA (Scharlab) at 22°C with a 16 h light/dark cycle. For long-term storage, *V. inaequalis* cellophane membranes were air-dried overnight and stored at –20°C. *Escherichia coli* strain DH5α (Thermo Fisher Scientific) was used for cloning, propagation and maintenance of Cas9HygAMA-*sgRNA* plasmids, and *E. coli* strain TG1 (kindly provided by Jasna Rakonjac, Massey University) for propagation of the Cas9HygAMAccdB plasmid, in lysogeny broth (LB) at 37°C and 180 rpm, or on LB agar at 37°C.

### Construction of the Cas9HygAMAccdB plasmid

The ANEp8_Cas9_LIC1 plasmid (35) was kindly provided by Concordia University and was adapted to contain a hygromycin cassette in place of the *pyrG* gene for selection. The 15.6-Kb ANEp8_Cas9_LIC1 plasmid was initially digested with the *NotI* restriction enzyme (New England Biolabs) to liberate a 5.3-Kb fragment containing the AMA1 cassette (purified by gel extraction using an Invitrogen PureLink Quick Gel Extraction Kit) and a 10.3-Kb fragment (purified by gel extraction) containing the *Cas9* and *pyrG* genes. Subsequent digestion of the 10.3-Kb fragment with the *KpnI* restriction enzyme (New England Biolabs) liberated a 9-Kb Cas9 cassette (purified by gel extraction) containing the *Cas9* gene and removal of *pyrG*. To amplify the hygromycin resistance cassette, PCR was performed on the plasmid pDONR221-Hyg (56) with the restriction enzyme-adapted primers KpnI TrpC Hyg DONR R and NotI PgpdA Hyg pDONR F (Table 2), and the resulting product was digested with *Kpn*I and *Not*I prior to ligation. The Cas9 (*Not*I/*Kpn*I-digested) cassette and the hygromycin resistance (*Not*I/*Kpn*I- digested) cassette were ligated with T4 ligase (Invitrogen) at 16°C overnight, creating the as9Hyg plasmid. The Cas9Hyg plasmid was re-digested with *NotI* and treated with alkaline phosphatase (purified by gel extraction) before its T4 ligation with the AMA1 (*Not*I-digested) cassette, creating the Cas9HygAMA plasmid. Modification to the sgRNA protospacer was achieved through ligation-independent cloning (LIC) (35). A mock sgRNA protospacer, containing two *Sap*I restriction enzyme sites, was cloned into the Cas9HygAMA plasmid, creating the Cas9HygAMASapI plasmid, and thus removing the need for the use of LIC with future protospacers. The primers required for the *Sap*I insertion using the LIC method were SapI site CRISPR Fw P1, SapI site CRISPR Rev P1, Fw LIC2 and Rev LIC2 (Table 2). A ccdB lethal cassette was cloned between the two *Sap*I (New England Biolabs) sites to aid in the efficiency of future protospacer cloning. To amplify the ccdB lethal cassette sequence (2 Kb), PCR was performed on the split marker vector pDONR-SM1 (57) with the *Sap*I restriction enzyme-adapted primers SapI ccdB F and SapI ccdB R (Table 2). The resulting product was digested with *Sap*I prior to its ligation with Cas9HygAMASapI (*Sap*I-digested), creating the Cas9HygAMAccdB plasmid.

### Protospacer design and cloning

The *V. inaequalis THN* gene was screened for CRISPR-Cas9 target sites with the PAM (NGG) sequence using Geneious v.9.0.5 software (58). Two protospacer sequences targeting the first and second exon of the *THN* gene, respectively, with the best on-target and off-target scores were selected. The selected protospacers were predicted to have no off-target binding sites in the *V. inaequalis* MNH120 PacBio reference genome (unpublished, The New Zealand Institute for Plant and Food Research Limited) using BLASTn. Each protospacer, with the appropriate *Sap*I overhang for Golden Gate cloning into the destination plasmid

Cas9HygAMAccdB, was ordered as a forward and reverse oligonucleotide from Integrated DNA Technologies. The protospacer #20 was generated by pre-annealing 40 ng of the forward (MR135) and reverse (MR136) oligonucleotides (Table 2), and protospacer #4 by pre-annealing 40 ng of the forward (MR137) and reverse (MR138) oligonucleotides (Table 2), in annealing buffer (10 mM Tris-HCl pH 8, 50 mM NaCl, 1 mM EDTA, pH 8) with the following thermocycler program: 5 min at 95°C, 20 sec at 92°C, followed by a decrease of 0.5°C each cycle for 140 cycles, and finally, 1 min at 25°C. Pre-annealed oligonucleotides were cloned into the Cas9HygAMAccdB plasmid using Golden Gate in association with the *Sap*I restriction enzyme to generate sgRNA20 and sgRNA4. Golden Gate reactions were performed with the following thermocycler program: 1 min at 37°C, 1 min at 16°C for 30 cycles, followed by 5 min at 55°C and 5 min at 80°C. Transformants positive for each Cas9HygAMA-*sgRNA* plasmid were screened by colony PCR using Taq DNA polymerase (New England Biolabs) with the forward Cas9HygAMAccdB primer (MR139) and reverse sgRNA-specific primer (MR136 or MR138) (Table 2). Colony PCRs were carried out with the standard manufacturer’s protocol. Sequence authenticity of the sgRNAs was confirmed by PCR amplicon sequencing, provided by the Massey Genome Service (Massey University, Palmerston North, New Zealand), using the MR139 forward Cas9HygAMAccdB primer.

### *V. inaequalis* protoplast preparation and transformation

Cas9HygAMA-*sgRNA* plasmids were introduced into *V. inaequalis* using a PEG-mediated protoplast transformation protocol. For this purpose, *V. inaequalis* was first grown on cellophane membranes overlaying PDA for 10–14 days. Fungal mycelia on top and inside cellophane membranes were then macerated in 1.5 ml microcentrifuge tubes using plastic micropestles, transferred to a 250 ml Erlenmeyer flask containing 30 ml potato-dextrose broth (PDB) (Difco^™^), and cultured without shaking in the dark at 22°C for 48 h. After culturing, fungal material was harvested by centrifugation at 2,800 *g* for 20 min, washed three times with KC buffer (0.60 M KCl, 50 mM CaCl_2_·2H_2_0), with collection by centrifugation as above after each wash, and incubated in a 250 ml Erlenmeyer flask containing 10 g/L *Trichoderma harzianum* lysing enzymes (Sigma-Aldrich) in 50 mL KC buffer at 24°C and 80 rpm for 4-5 h. Finally, protoplasts were filtered through glass wool, and washed three times with KC buffer as above. Protoplasts were counted using a haemocytometer and re-suspended to a final concentration of 10^4^–10^5^ protoplasts/ml.

Transformation was performed by mixing 100 μl of *V. inaequalis* protoplasts (10^4^-10^5^ protoplasts/ml) with 100 μl of 25% PEG4000, 10 μg of circular *Cas9HygAMA-sgRNA* plasmid DNA, and 5 μl of 50 μM sterile spermidine. The protoplast-PEG mixture was then chilled on ice for 20 min and 500 μl of 25% PEG4000 gently added. Each protoplast-PEG mixture was plated across five Sucrose Hepes (SH) plates (0.6 M sucrose, 5 mM HEPES, 0.6% agar) and incubated at 20°C for 48 h. After incubation, protoplasts were overlaid with ½-strength PDA cooled-down to ~50°C and supplemented with 50 mg/ml hygromycin B (Merck). Transformants appearing on the PDA surface, between two and three weeks after transformation, were transferred to 16-well PDA plates supplemented with 50 mg/ml hygromycin B, and grown until abundantly sporulating. After mutant screening, selected transformants were single-spore purified. This was achieved by re-suspending a single colony in 500 μl sterile water and vortexing for 30 sec, with 100 μl streaked onto 4% water agar (WA) plates and the conidia germinated for 24 h. Following germination, one single germinated conidium was transferred to a cellophane membrane overlaying PDA for continued growth.

### *V. inaequalis* genomic DNA extraction

Two-to-four week-old cultures of *V. inaequalis* grown on cellophane membranes were freeze-dried and ground to a fine powder in liquid nitrogen with a pre-cooled mortar and pestle, and approximately 300 mg of powder was transferred to a 1.5 ml microcentrifuge tube. To this, 1 ml of DNA extraction buffer (0.5 M NaCl, 10 mM Tris-HCl, 10 mM EDTA, 1% SDS, pH 7.5) was added, vortexed, and incubated at 65°C for 30 min, followed by 2 min incubation at room temperature (RT). Fungal material was collected by centrifugation at 16,000 *g* for 2 min and 800 μl of supernatant was transferred to a fresh 1.5 ml microcentrifuge tube. Then, 4 μl of RNase A (20 mg/ml) (Invitrogen) was added and samples were incubated at 37°C for 15 min. After incubation, a 0.5 volume of phenol and a 0.5 volume of chloroform:isoamyl alcohol (24:1) were added and samples were centrifuged for 5 min at 16,000 *g*. The aqueous phase was then transferred to a fresh 1.5 ml microcentrifuge tube and 1 volume of phenol:chloroform (1:1) was added. Samples were again centrifuged at 16,000 *g* for 5 min and the supernatant transferred to a new 1.5 ml microcentrifuge tube. The chloroform:isoamyl alcohol (24:1) step was then repeated and the supernatant was transferred to a new 1.5 ml microcentrifuge tube. Genomic DNA was precipitated by the addition of a 0.1 volume of 3 M sodium acetate (pH 5.2) and two volumes of 95% ethanol. Samples were mixed by inversion and incubated overnight at –20°C. Following incubation, the precipitated DNA was collected by centrifugation at 16,000 *g* for 30 min, and the supernatant was decanted. The genomic DNA pellet was then washed with 200 μl of 70% ethanol and collected by centrifugation at 16,000 *g* for 5 min. Finally, the genomic DNA pellet was air-dried for 15-30 min and suspended in 50 μl of MilliQ water.

### High resolution melting curve analysis

A positive control for the HRM curve analysis was created by site-directed mutagenesis. First, primers MR161 and MR162 (Table 2) were phosphorylated with T4 Polynucleotide Kinase (New England Biolabs) at 37°C for 1 h in a 4.5 μl reaction volume with 10 μM primer, 10x T4 ligase buffer (New England Biolabs) and 0.4 μl of T4 polynucleotide kinase (New England Biolabs). The *THN* gene was amplified with the phosphorylated primers MR161-MR162 using Phusion Flash High-Fidelity PCR Master Mix (Thermo Fisher Scientific) and purified using an OMEGA Gel Extraction kit. The pICH41021 plasmid was digested with the *Sma*I restriction enzyme in a 50 μl volume for 2 h at 37°C. Digested plasmid was then de-phosphorylated with Shrimp Alkaline Phosphatase (rSAP) (New England Biolabs) for 30 min at 37°C, with the reaction subsequently heat-inactivated at 65°C for 5 min. Finally, the *THN* gene was ligated to pICH41021 in a 3:1 molar ratio using T4 Ligase (New England Biolabs) at 4°C overnight and transformed by heat shock into chemically competent *E. coli* cells. Positive transformants were confirmed by colony PCR. Next, the pICH41021-*THN* plasmid was amplified with overlapping primers MR182 and MR183 to introduce a mutation in the *THN* gene that substituted a thymine and cytosine at nucleotide positions 26 and 27 for two guanines. The resulting re-circularized pICH41021-THN_TC(26/27)GG_ plasmid was confirmed by sequencing.

Two sets of primers (MR170-MR171 and MR172-MR173) were designed with Geneious v.9.0.5 software for use in the HRM curve analysis (Table 2). These primers were designed to amplify the DNA region in the *THN* gene recognized by the sgRNA (and thus, edited by the Cas9 endonuclease), with an amplicon size of 123 bp (MR172-MR173) and 230 bp (MR170-MR171). Primers were tested to be specific and suitable for HRM by performing an HRM curve analysis with DNA standards (wt and engineered positive control), as described below, with the resulting amplicons resolved by electrophoresis on a 1.5% TAE gel.

The HRM curve analysis was performed using a LightCycler 480 Instrument (Roche) with the AccuMelt HRM SuperMix fluorescent dye (DNAture) in a 20 μL reaction with 1x AccuMelt HRM SuperMix, 300 nM forward primers (MR172), 300 nM reverse primer (MR173) and 1.5 ng of genomic DNA template or 0.01 ng of the puc19-THN_TC(26/27)GG_ plasmid. At least two technical replicates were performed for each sample. The qPCR amplification was performed with the following program: initial denaturation of 5 min at 95°C, followed by 40 cycles of 95°C for 8 sec, 60°C for 15 sec, 70°C for 20 sec, with one fluorescence reading per annealing step. The qPCR was followed by a melting program consisting of 95°C for 1 min, 40°C for 1 min, 76°C for 1 sec, and then a continuous 92°C with 25 acquisitions per degree followed by a cooling step of 40°C for 30 sec. The HRM curve data were analysed with the LightCycler 480 gene scanning software. To confirm mutants identified by HRM, the *THN* gene was amplified with Phusion Flash High-Fidelity PCR Master Mix using primers MR161 and MR162, with the resultant PCR amplicons gel-purified as above and sequenced using primer MR161.

### Ethics approval and consent to participate

Not applicable.

## Availability of data and materials

No substantive datasets were generated during the course of this study and all data are presented within.

## Author contributions

MR, JKB, KMP, REB, RDJ, LJJ and CHM conceived the project. MR, SA, DH and JS performed the research. All authors contributed to the writing and reviewing of the manuscript.

## Acknowledgements

We thank Professor Adrian Tsang (Center for Functional and Structural Genomics Biology, Concordia University, Canada) for hosting JS and providing the original CRISPR-Cas9 vectors. We also thank Natasha Forester and Pranav Chettri for useful discussions around vector modification.

