## Supplementary material for "CRISPR-Cas9 gene editing and rapid detection of gene-edited mutants using high-resolution melting in the apple scab fungus, *Venturia inaequalis*": Figure S1

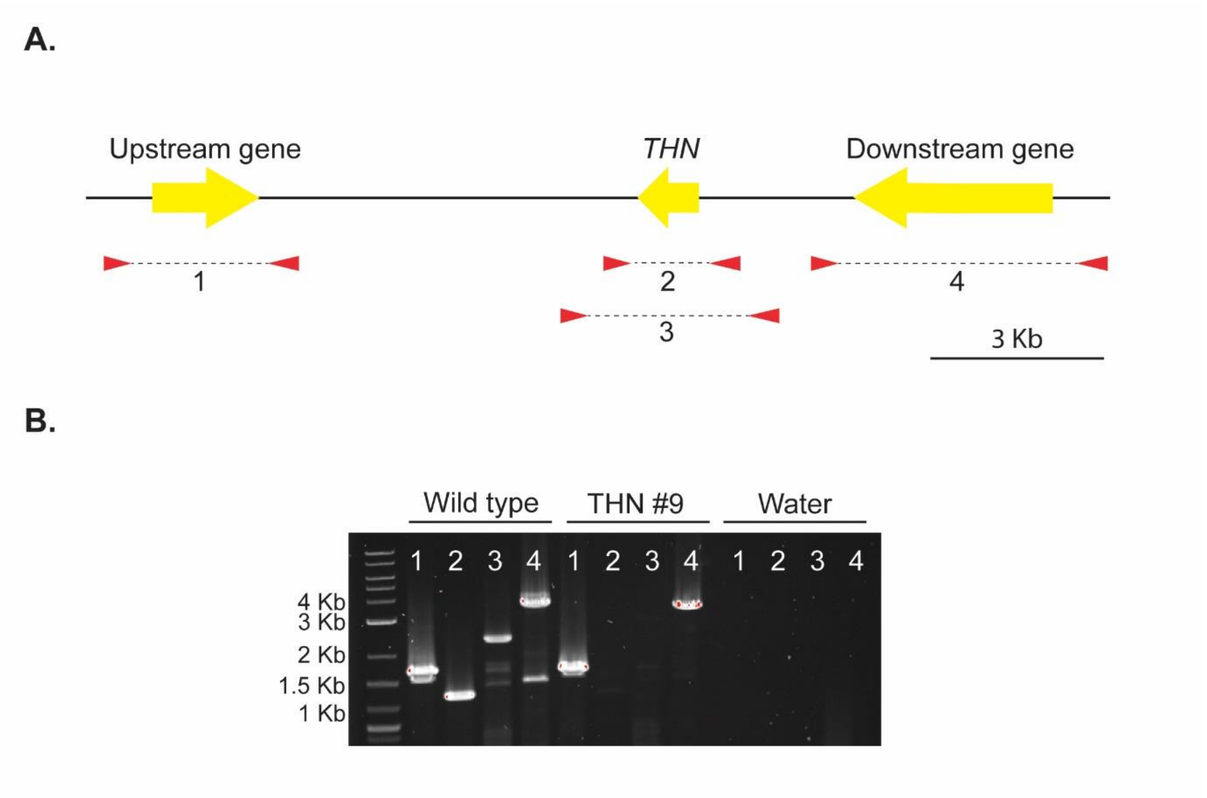


**Figure S1. A large deletion event has potentially occurred at the *THN* locus in mutant THN #9. A.** Schematic representation of the *trihydroxynaphthalene reductase* (*THN*) gene locus in the *Venturia inaequalis* MNH120 genome. Arrows: primer binding site, dashed lines: amplicon. **B.** Conventional PCR of *THN* and its neighbouring genes using wild type genomic DNA, THN #9 genomic DNA and water as a negative control. PCR amplicons were resolved by electrophoresis on a 1% TBE gel. Marker: 1 Kb plus DNA ladder. 1. MR204-MR205: upstream gene; 2. MR161-MR162: *THN*; 3. MR184-MR185: *THN*; 4. MR252-MR253: downstream gene.
