## Supplementary material for "CRISPR-Cas9 gene editing and rapid detection of gene-edited mutants using high-resolution melting in the apple scab fungus, *Venturia inaequalis*": Figure S2

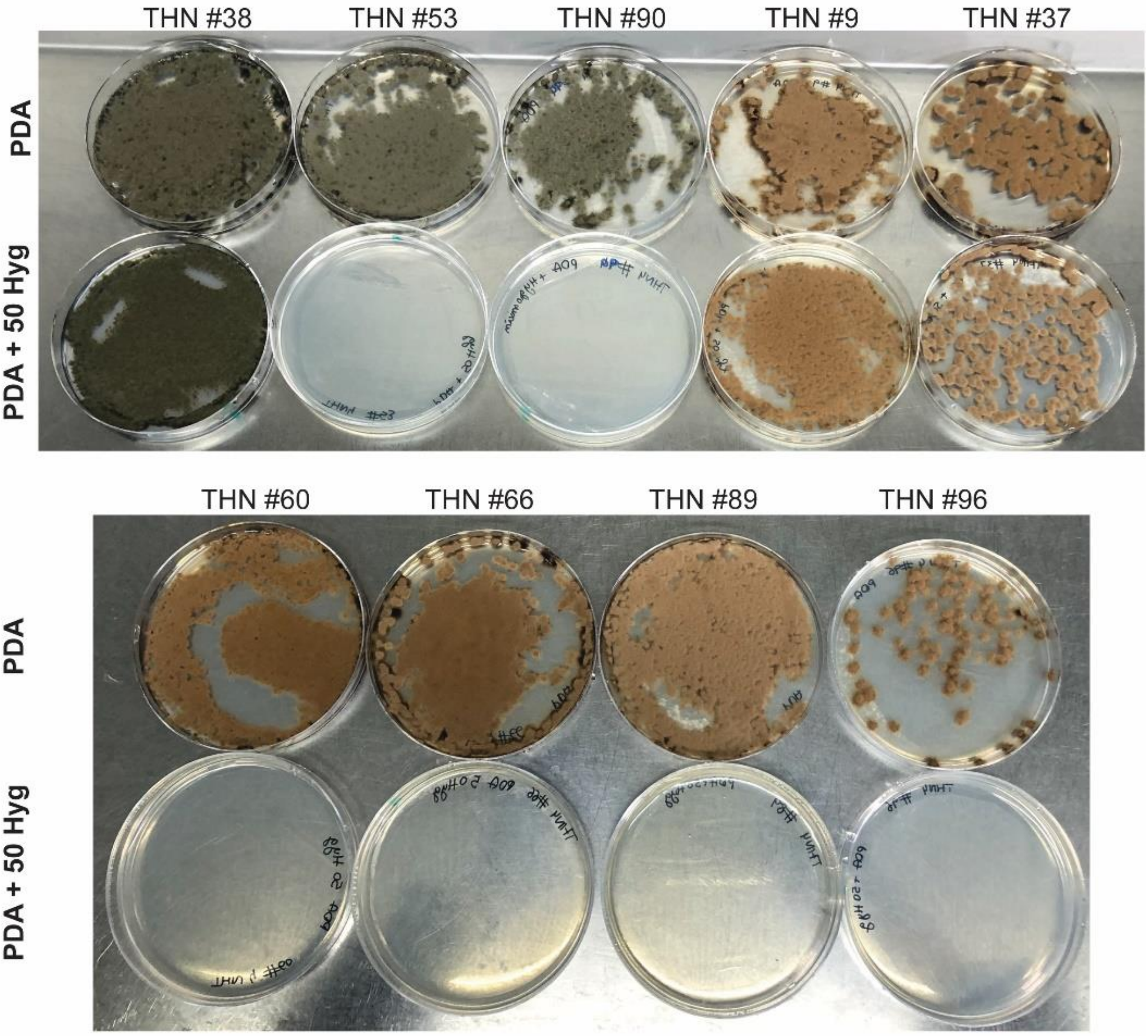


**Figure S2. Loss of the autonomously replicating Cas9HygAMA-*sgRNA* plasmid by CRISPR-Cas9 *THN* transformants of *Venturia inaequalis*.** Conidia of each transformant were plated on a cellophane membrane overlaying potato-dextrose agar (PDA) or a cellophane membrane overlaying PDA supplemented with 50 µg/ml hygromycin B (Hyg), followed by incubation at 22°C for 14 days.
